# Shifts in Coral Reef Holobiont Communities in the High-CO_2_ Marine Environment of Iōtorishima Island

**DOI:** 10.1101/2024.12.02.626509

**Authors:** Roger Huerlimann, Hin Boo Wee, Maria Alves dos Santos, Hiroki Kise, Masaru Mizuyama, ‘Ale’alani Dudoit, Emmeline Jamodiong, Nagi Satoh, Giun Yee Soong, Haruko Kurihara, Robert J Toonen, Filip Husnik, Akira Iguchi, Timothy Ravasi, James Davis Reimer

## Abstract

Ocean acidification (OA), driven by rising atmospheric CO_2_, presents a serious threat to marine biodiversity, especially within coral reef ecosystems. Natural analogue sites, such as the high-pCO2 seep at Iōtorishima Island in Japan, offer insights into future conditions. This study investigated the holobiont communities of Symbiodiniaceae and bacteria in the zoantharian *Palythoa tuberculosa* at Iōtorishima and compared them to specimens from control sites in Okinawa and Hawaiʻi.

Using amplicon sequencing of the dinoflagellate internal transcribed spacer 2 (ITS2) region of ribosomal DNA and microbial 16S rRNA gene, we detected significant shifts in both Symbiodiniaceae and bacterial communities under high-pCO_2_ conditions at Iōtorishima. Specifically, *P. tuberculosa* at the seep site had reduced Symbiodiniaceae diversity, predominatly featuring *Cladocopium* C1 and C3 types. Additionally, its bacterial communities showed lower richness with distinct taxonomic profiles, including increased levels of *Mollicutes* and *Vibrio* spp.

These results highlight the potentially adverse effects of OA on hexacoral holobionts and emphasize the need for detailed, high-resolution studies across various holobiont species and geographic locations. The shifts observed specifically in Symbiodiniaceae and bacterial communities at the Iōtorishima Seep suggest that holobionts may exhibit plasticity in response to environmental stress, which has implications for resilience and adaptation of zoantharians and other reef organisms amid climate change. This research provides crucial baseline data for predicting future coral reef compositions in an OA-affected world.

## Introduction

Under climate change, future oceans are expected to have higher pCO_2_ levels and lower pH compared to today. Over the past thirty years, numerous studies have documented a wide range of potential negative impacts of these changes on various marine organisms (e.g.; (Havenhand, Buttler, Thorndyke, & Williamson, 2008; Heuer & Grosell, 2014; Hoegh-Guldberg et al., 2007; Rodolfo-Metalpa et al., 2011; Rodolfo-Metalpa, Lombardi, Cocito, Hall-Spencer, & Gambi, 2010). However, significant knowledge gaps remain, and a comprehensive understanding of future oceans ecosystem is still a subject of debate (Leung, Zhang, & Connell, 2022). One promising approach to improve our predictions of future changes in marine ecosystem is the study of natural analogues.

Natural analogues of future oceans are typically areas where current environmental conditions resemble those projected for the future. These sites include locations like CO_2_ seeps and enclosed bays where limited water exchange allows lower pH and higher pCO_2_ conditions, often due to decaying biomass, to persist (Agostini et al., 2021; Rodolfo-Metalpa et al., 2010). Notable examples of these natural analogue sites include Vulcano Island in Italy (Boatta et al., 2013), Nikko Bay in Palau (Golbuu, Gouezo, Kurihara, Rehm, & Wolanski, 2016) and Ambitle in Papua New Guinea (Pichler et al., 2019).

Shallow water subtropical and tropical coral reef ecosystems harbor the highest levels of marine biodiversity (Reaka-Kudla, 1997), primarily constructed by the skeletal secretions of organisms, such as zooxanthellate scleractinian corals, coralline algae, and foraminifera (e.g., Jindrich (1983)). These bioengineers, along with their calcium carbonate and aragonite skeletons, may be particularly susceptible to increased pCO_2_ concentrations and decreased pH levels (Hoegh-Guldberg et al., 2007). Therefore, research at natural analogue sites within coral reefs in subtropical and tropical regions especially crucial for gaining knowledge to better protect marine biodiversity in the face of future environmental conditions.

Research on subtropical and subtropical natural analogue sites has been relatively limited so far, and the findings vary depending on the species and study (Leung et al., 2022). Benthic studies have primarily focused on scleractinian corals (Leung et al., 2022), which is understandable given their key role as ecosystem engineers. Some studies have reported a general decline in scleractinians, accompanied by increases in other anthozoans such as soft corals (Inoue, Kayanne, Yamamoto, & Kurihara, 2013), zoantharians (Reimer et al., 2023), and sea anemones (Suggett et al., 2012). However, these other groups have not been investigated as extensively as scleractinians. Research on the photosymbionts of these anthozoans, the Symbiodiniaceae dinoflagellates (LaJeunesse et al., 2018), has produced contrasting results. While some studies found that Symbiodiniaceae diversity remained unaffected by natural analogue conditions (e.g., in scleractinians; (Noonan, Fabricius, & Humphrey, 2013), others have indicated negative impacts on diversity (Wee, Kurihara, & Reimer, 2019). Additionally, there have been very few studies on other holobiont components such as bacteria, protists, or viruses at natural analogues (e.g., (Yang et al., 2020). Even in existing research, methods are often basic, with limited resolution and scope. It is evident that there is a significant data gap, highlighting the urgent need for advanced, large-scale, and multi-faceted datasets focusing on non-scleractinian benthos from coral reef natural analogues, especially given the apparent resilience of some of these groups to low pH conditions (Inoue et al., 2013; Reimer et al., 2023; Suggett et al., 2012).

One such natural analogue site is located at the uninhabited Iōtorishima Island in the Ryukyu Islands of southern Japan. The Iōtorishima site features a CO_2_ seep over a shallow coral reef (Inoue et al., 2013), where environmental conditions within the inner lagoon vary based on seep activity, seasons, and tides, and sometimes leading to high pCO_2_ levels (Inoue et al., 2013). Initial studies at Iōtorishima revealed an increase in soft coral abundance (Inoue et al., 2013) at the high-CO_2_ site, and more recent research has also documented numerous zoantharians (Reimer, Wee, López, Beger, & Cruz, 2021) and the “living fossil” octocoral *Nanipora* (Reimer, Kurihara, et al., 2021), related to blue corals. These finding suggests that understudied non-scleractinian groups may often thrive in marginal and unique environments (Reimer et al., 2023; Suggett et al., 2012), indicating a need for closer investigation of these taxa.

In this study, we used high-throughput sequencing to identify the Symbiodiniaceae and bacteria communities of the zoantharian *Palythoa tuberculosa* at Iōtorishima, Okinawa Island, and Hawai’i. By comparing the composition of microbes associated with *P. tuberculosa* across these regions, we gain a robust image of how holobionts change under low pH and high pCO_2_ levels.

## Materials and Methods

### Iōtorishima site and specimen collection

Iōtorishima Island (hereafter Iōtorishima) is a small uninhabited island in the middle Ryukyu Islands in the East China Sea. The island is approximately 65 km to the west of Tokunoshima Island of Kagoshima, and is the northernmost island in Okinawa Prefecture. Iōtorishima was inhabited until 1958 when volcanic activity forced the final evacuation of residents. In recent years, despite being isolated and uninhabited, the island has been the focus of marine scientific research detailing a natural analogue as there is a CO_2_ vent in a shallow fringing reef on the southeast coast of the island (Inoue et al., 2013; Reimer, Kurihara, et al., 2021; Reimer, Wee, et al., 2021; Wee et al., 2019).

As detailed in (Reimer, Kurihara, et al., 2021), Iōtorishima was visited from 14. to 16. September in 2020 (Fig. 1, Table 1). Field surveys focused on the shallow CO_2_ seep at depths of 0.5 to 2 m within the shallow coral reef lagoon (Inoue et al., 2013), centered around 27° 52′ 11.8″ N, 128° 14′ 01.8″ E. During benthic surveys, *Palythoa tuberculosa* (Esper, 1805) specimens (n = 14) were collected within the shallow CO_2_ lagoon by snorkeling (=high CO_2_). We also collected specimens of the same species from a control site (n = 7) within the lagoon at the same depths with lower CO_2_ levels; this was the same as described in (Wee et al., 2019). *P. tuberculosa* is easily identifiable in the field by its ‘immersae’ colony morphology (Pax, 1910; Reimer, 2010). Tissue samples were individually preserved in >95% ethanol and stored at −20°C until further analyses.

**Fig. 1.**
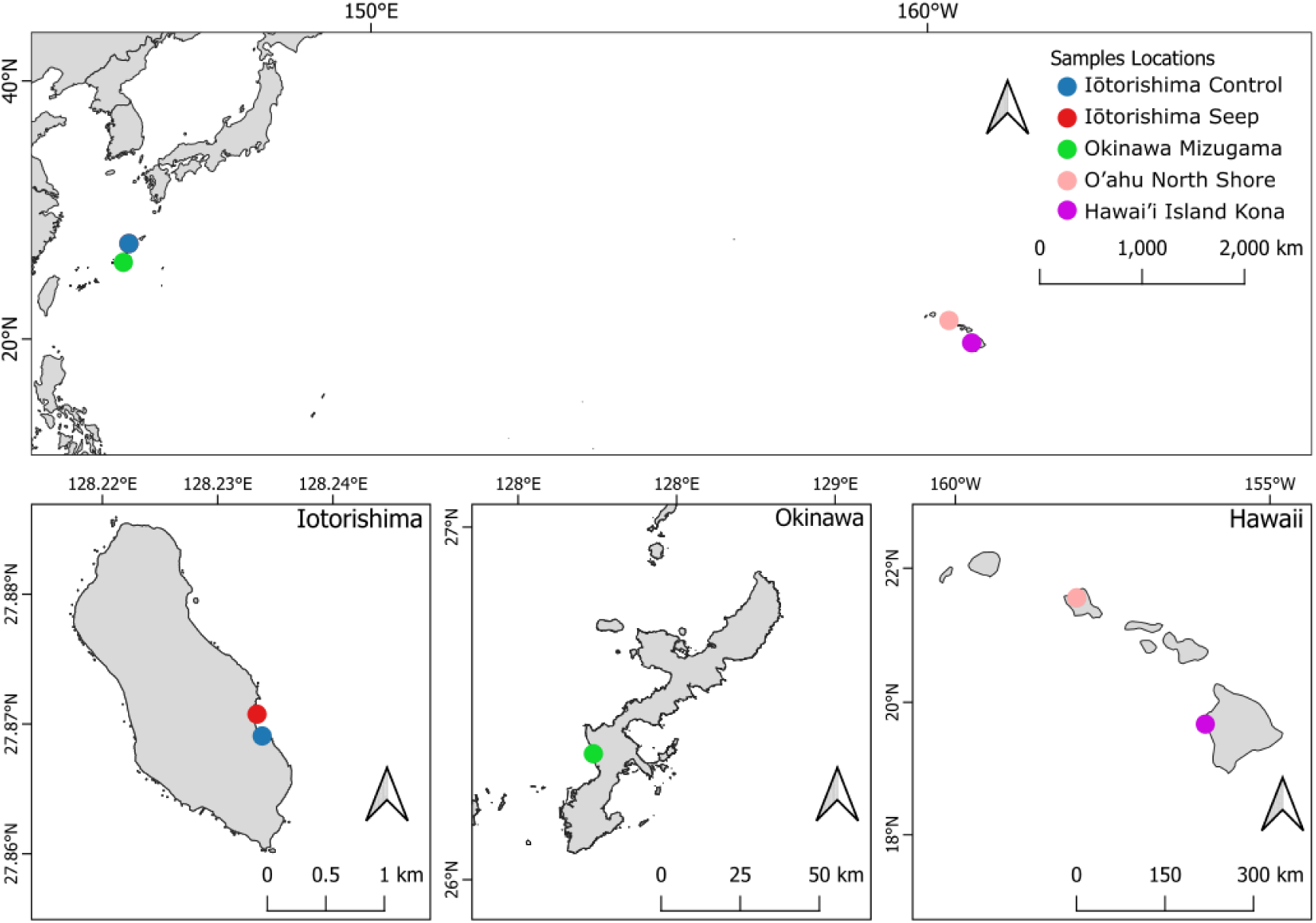
Locations of sampling sites of *Palythoa tuberculosa* examined in this study across the northern Pacific Ocean.

**Table 1.** Sampling details.

| Location | Sample Numbers | Date Collected | Depth | Coordinates |
| --- | --- | --- | --- | --- |
| Iōtorishima Control | 7 | 14. to 16. Sep. 2020 | 0.5 to 2 m | 27°52'09.8"N<br>128°14'04.1"E |
| Iōtorishima Seep | 14 | 14. to 16. Sep. 2020 | 0.5 to 2 m | 27°52'15.1"N<br>128°14'02.0"E |
| Okinawa Mizugama | 29 | 13: 17. Aug. 2017 and<br>16: 9. Apr. 2018 | 0.5 to 10 m | Specimen 1 to 5:<br>26.358164°<br>N,127.737515° E<br>Specimen 6 to 10:<br>26.358255°<br>N,127.737834° E<br>Specimen 11 to 15:<br>26.360122°<br>N,127.739596° E<br>Specimen 16 to 20:<br>26.355274°<br>N,127.739291° E<br>Specimen 21 to 25:<br>26.355656°<br>N,127.740174° E |
| O'ahu North Shore | 16 | 10. Sept. 2020 | 1 to 2m | 21.561657° N,<br>158.071598° W |
| Hawai'i Island Kona | 9 | 20. July. 2018 | 1 to 2m | 19.6719° N,<br>156.0269° W |

### Mizugama (Okinawa) site

*Palythoa tuberculosa* samples were collected on 17th August 2017 (n = 13) and 9th April 2018 (n = 16 specimens) at the Mizugama seawall reef (Fig. 1, Table 1). The specimens were collected via SCUBA diving or reef walking at low tide. Specimens were collected at different distances from the Mizugama river mouth and at various depths: specimens No. 1-5 (0 m from river mouth, 10 m depth), No. 6-10 (0 m from river mouth, 2 m depth), No. 11-15 (tidal pool inside river mouth, 0.5 m depth), No. 16-20 (500 m from river mouth, 10 m depth), No. 21-25 (500 m from river mouth, 2 m depth). Collected specimens were preserved in ethanol 99% and transferred to the laboratory and stored at −20°C until subsequent analyses.

### Hawaiian Islands sites

We sampled 25 individuals of *Palythoa tuberculosa* from the north shore of Oʻahu and Kona coast of Hawaiʻi Island (Fig. 1, Table 1) between July 2018 to September 2020 at depths of 1 to 2 m. Collections were made via snorkeling or SCUBA diving, and tissue samples were individually preserved in salt-saturated DMSO (dimethyl sulfoxide) buffer and stored at room temperature (Gaither, Szabó, Crepeau, Bird, & Toonen, 2011) until further analyses.

### DNA extraction and high-throughput sequencing

Genomic DNA of *Palythoa tuberculosa* collections was extracted using the Qiagen DNEasy PowerSoil kit (Tokyo, Japan) following the manufacturer’s protocols. The concentration of the genomic DNA samples was analyzed using a Qubit Fluorometer with the Qubit 1X dsDNA HS Assay kit (Invitrogen, Japan).

For Symbiodiniaceae, amplicon libraries were generated from the internal transcribed spacer 2 region of ribosomal DNA (ITS2 rDNA). The first PCR was performed with the primer set SYM_VAR_5.8S2/SYM_VAR_REV (Hume et al., 2018) with an overhang adapter sequence for the MiSeq platform (Illumina, Los Angeles, CA, USA). Each 20 µl PCR included the following components: 1.0 µl of temperate DNA, 1.2 µl of each forward and reverse primer, 0.2 µl of TaKaRa Ex Taq™, 2.0 µl of 10×Ex Taq Buffer, 1.6 µl of dNTP mixture (Takara Bio Inc., Shiga, Japan), and 12.8 µl of nuclease-free water. The PCRs were run with a temperature profile of 98°C for 2 min, followed by 35 cycles of 98 °C for 10 s, 56°C for 30 s, and 72°C for 30 s, with a final extension at 72 °C for 7 min. Generated amplicons were purified using Ampure XP beads (Beckman Coulter, Brea, CA, USA). Index PCR was performed to add Illumina sequencing adaptors and sample index sequences, followed by second purification using Ampure XP beads. Quantification of the concentration of purified index PCR products was measured by the Qubit dsDNA HS Assay Kit (Thermo Fisher Scientific, Waltham, MA, USA). These PCR products were pooled at equimolar concentrations. Amplified PCR products were sequenced on an Illumina MiSeq platform at the National Institute of Advanced Industrial Science and Technology, using a V2-500 cycle kit to generate 2 × 250 bp paired-end reads.

For bacteria, we amplified the V3-V4 region of the bacterial 16S rRNA gene using the primer set P341F/P805R (Takahashi, Tomita, Nishioka, Hisada, & Nishijima, 2014) with overhang adaptor sequences. Each 20 µl PCR included the following components: 1.0 µl of temperate DNA, 0.4 µl of each forward and reverse primer, 0.4 µl of MightyAmp DNA Polymerase, 10 µl of 2x MightyAmp Buffer (Takara Bio Inc., Shiga, Japan), and 7.8 µl of nuclease-free water. The PCRs were run with a temperature profile of 98 °C for 2 min, followed by 10 cycles of 98 °C for 10 s, 65–56°C for 15s (decrease temperature by 1°C per cycle) and 68°C for 1 min, and 25 cycles of 98 °C for 10 s, 55°C for 15s and 68°C for 1 min, with a final extension at 68°C for 2 min. Index PCR was performed to add Illumina sequencing adaptors and sample index sequences. Index PCR products were purified using Ampure XP beads and quantified using the Qubit dsDNA HS Assay Kit, followed by pooling at equimolar concentrations. The purified library was sequenced on an Illumina Miseq platform, using a V3-600 cycle kit to generate 2 × 300 bp paired-end reads.

### Bioinformatical analysis

For Symbiodiniaceae, the demultiplexed paired-end reads were analyzed in local using SymPortal analytical framework (Hume et al., 2019), a platform for phylogenetically resolving Symbiodiniaceae taxa using ITS2 amplicon data. Sequence quality control (QC) of paired-end reads was performed utilizing mothur 1.39.5 (Schloss et al., 2009), blast+ suite of executables (Camacho et al., 2009) and minimum entropy decomposition (Eren et al., 2015) to filter artifactual and non-Symbiodiniaceae sequences from dataset. Post QC sequences from each sample were loaded into the Symportal database to identify the specific sets of sequences called defining intragenomic variants (DIVs).

For bacteria, the demultiplexed paired-end sequence reads were analyzed in the QIIME2 v2022.8.3 framework (Bolyen et al., 2019), using several plug-in programs (Hamamoto et al., 2024). End sequences, including primer and adopter sequences, were trimmed from the 5’ and 3’ ends using Cutadapt v4.1 (Martin, 2011) (via q2-cutadapt) for up to 10 repeats. If the primer sequences were not found or the trimmed length of the sequences was less than 100 bp, the reads were discarded. Quality control, including quality filtering, denoising, correction of reading errors, merging of paired-end reads, and removal of non-biological sequences (including chimeras), was performed by DADA2 v4.1.3 (Callahan et al., 2016) (via q2-dada2) with the following custom values. For denoising, the maximum expected error values, ‘maxEE’ (Edgar & Flyvbjerg, 2015), were set to 2 for forward reads and 5 for reverse reads. The truncated length at the 3’ end of forward and reverse reads was independently determined by the base position corresponding to a Phred quality score of less than 20 in the first quartile of total reads. After quality control, a count table of representative sequences was generated by dereplication of the same amplicon sequence variant (ASV).

Taxonomic assignment to the ASV was performed using a pre-trained Naive Bayes classifier (Bokulich et al., 2018) with reference to the 16S ribosomal RNA sequence database SILVA v138.1 (Quast et al., 2012; Yilmaz et al., 2014) curated by RESCRIPt (Robeson et al., 2021), and then operational taxonomic units (OTUs) were determined for each ASV using scikit-learn (Pedregosa et al., 2011), a machine learning classifier plugin in Qiime2.

The taxonomic assignment identified one genus as “Candidatus *Hepatoplasma*”; however, in our experience this is a misidentification due to the uncertainty within this group of marine bacteria. The alignment of our ASV sequences against sequences within this group showed a close relationship with the poorly research clade *Metamycoplasmataceae* (Vohsen et al., 2024). Considering the importance of this group in our findings, coupled with the uncertainty in the taxonomy, we decided to rename the ASVs as “Unknown Genus within *Mollicutes*” or in places “*Mollicutes*” for short (see Supplemental Figure 1). Future work will hopefully further resolve this important bacterial group.

The average number of reads per sample was 44,344 ± 26,807 (mean ± stdev) before filtering, and 38,181 ± 26,535 (mean ± stdev) after filtering. One sample each from Okinawa Mizugama and Hawai’i Kona were removed due to low read counts (Mizugama: 1,856 reads, Kona: 136 reads) and distant clustering from other related samples. Other filtering included removing ASVs (amplicon sequence variants) with no taxonomic assignment at the Phylum level (i.e., “NA”), filtering ASV assigned to Chloroplast at the Order level, Mitochondria at the Family level, and *Cutibacterium* at the Genus level.

### Statistical analyses

The NGS reads of family-level taxonomic clustering of bacteria and genus-level clustering of Symbiodiniaceae for *Palythoa tuberculosa* were optimised and organised based on ASV and DIV, respectively. One table each was produced for bacteria and Symbiodiniaceae data, and the reads were transformed into percentage abundance of composition for each specimen/individual. The specimens were categorized based on the locations they were collected: Iōtorishima (seep), and coral reef sites with standard *pH* levels of Iōtorishimal (control), Okinawa (Mizugama), and Hawai’i (Kona, Hawai’i Island and North Shore, O’ahu).

Both bacteria (Family, Species, and ASVs) and Symbiodiniaceae datasets were analyzed using R (version 4.3.2; (R Core Team, 2013) via RStudio V2023.06.1 Build 524 (Racine, 2012)(RStudio, 2023). Alpha-diversity was analyzed using the phyloseq package (McMurdie & Holmes, 2013).

Ordinated (Bray-Curtis Dissimilarity) tests of beta-diversity dispersion and distribution (permutest) based on locations were conducted for both datasets. Permutative ANOVA (PERMANOVA) analyses via adonis2 (vegan package, version 2.6.4, (Dixon, 2003; Oksanen et al., 2013)) were conducted to test the dissimilar distance of composition among the sites, followed by a post-hoc test (pairwiseAdonis package). Similarity Profiling (SIMPER) analyses were conducted to test the apriori grouping of the compositions via clustering. The distance differences among the specimens and locations were represented in Non-parametric multidimensional scaling (nMDS).

Figures were created via R (base and ggplot2 package (Wickham, 2011)). Post-hoc tests were conducted with Holm’s correction method. The alpha value of all the statistical tests were set at α=0.05. Both alpha diversity measures were significant for Shapiro-Wilk normality test, therefore the non-parametric Kruskal-Wallis rank sum test and pairwise comparisons using Wilcoxon rank sum exact test were used for the statistical analysis.

## Results

### Alpha diversity - Bacteria

In terms of alpha diversity, there was a significant difference (Kruskal-Wallis chi-squared = 28.2, df = 4, p-value = 1.115e^-05^) in the bacterial richness at the ASV level between all locations. A post-hoc comparison showed that the Iōtorishima Seep location had a significantly lower richness than any other site (Fig. 2). In contrast, the post-hoc analysis of the evenness showed no significant differences between locations (Fig. 2), despite an overall significant result (Kruskal-Wallis chi-squared = 13.0, df = 4, p-value = 0.01124).

**Fig. 2.**
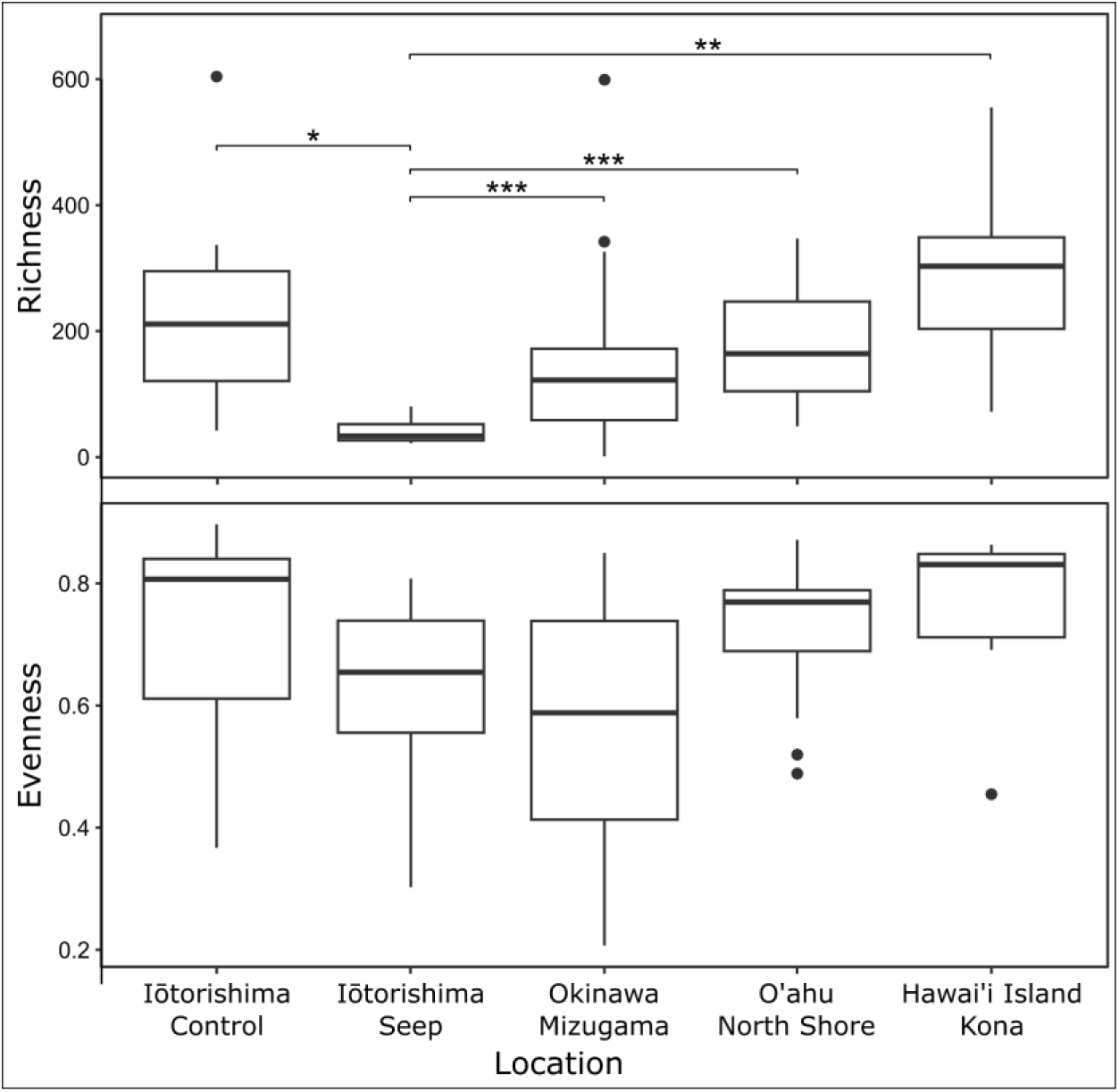
Microbial richness and evenness at the ASV level of the five locations. (statistical significance: * = 0.05, ** = 0.01, *** = 0.001; outliers are marked with filled circles)

### Beta diversity - Bacteria

For the bacterial beta diversity, there were significant differences among the locations regarding the family composition of bacteria hosted by *Palythoa tuberculosa* (PERMANOVA: R^2^=0.204, F= 4.032, p<0.001). Tests of composition dissimilarity between groups (with Holm’s correction) showed only Hawai’i Island Kona recorded no significant differences in bacterial family composition with Iōtorishima Control and O’ahu North Shore (Table 2A, pairwise Adonis: p = 0.356). The species composition of bacteria in *P. tuberculosa* was significantly different among the locations (PERMANOVA: R^2^=0.196, F=3.831, p<0.001). Pairwise tests showed only Hawai’i Island Kona and Okinawa Mizugama bacterial species compositions were not significantly different (Table 2B, pairwise Adonis: R^2^= 0.053, p = 0.053). Lastly, the highest resolution was found at the ASV level, which showed significant differences among (PERMANOVA: R^2^=0.164, F=3.0872, p<0.001) and between locations (Table 2C, pairwise Adonis: p < 0.010).

**Table 2.**
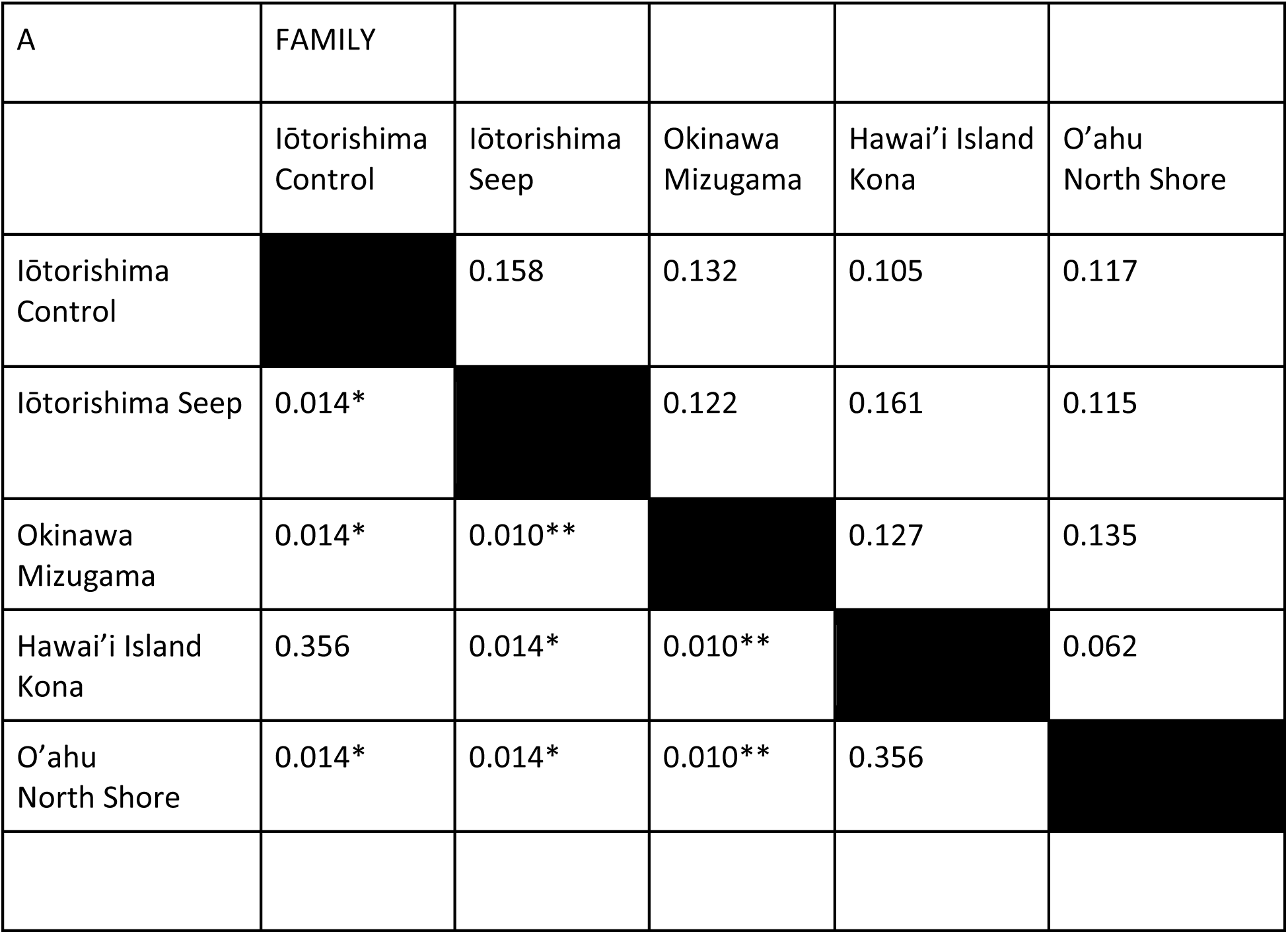
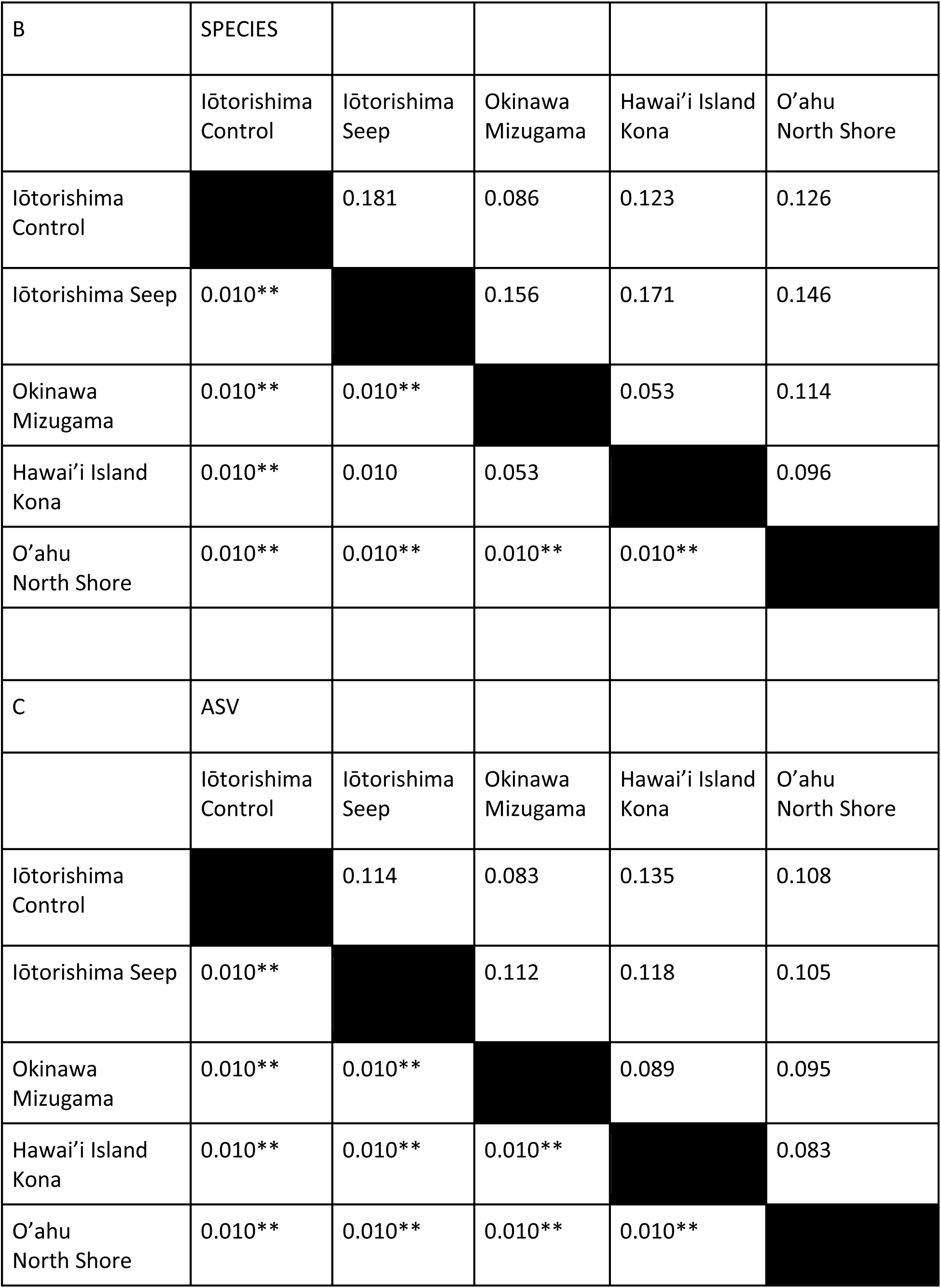
Pairwise PERMANOVA (Pairwise Adonis) conducted on the dissimilarity of bacteria composition (OTU: A=Family, B = Species, C =ASV) hosted by *Palythoa tuberculosa* for different locations. The lower F represents the adjusted p-values (Holm’s correction) and the upper table represents the R2 values. (statistical significance: * = 0.05, ** = 0.01)

nMDS showed more defined separations of dissimilarities among the locations as the OTU resolution increased from family (Figure 3A, stress=0.204) to ASV (Figure 3E, stress=0.216). Iōtorishima Seeps were observed to have more dissimilarities with Iōtorishima Control and Hawai’i Island Kona for species (Figure 3C, stress=0.219); and ASVs saw the same separation among the locations in addition to statistically significant differences with Okinawa Mizugama. This nMDS also showed that the site closest to Iōtorishima Seeps was O’ahu North Shore. nMDS graphs clearly showed that the Iōtorishima Seep had the most distinct composition among all the locations in the study. However, there were no significant beta-dispersions of the diversity among the locations regarding family (Permutest: Df=4, F=0.812, p=0.519), species (Permutest: Df=4, F=0.847,p=0.479), or ASV (Permutest: Df=4, F=0.315,p=0.858).

**Fig. 3.**
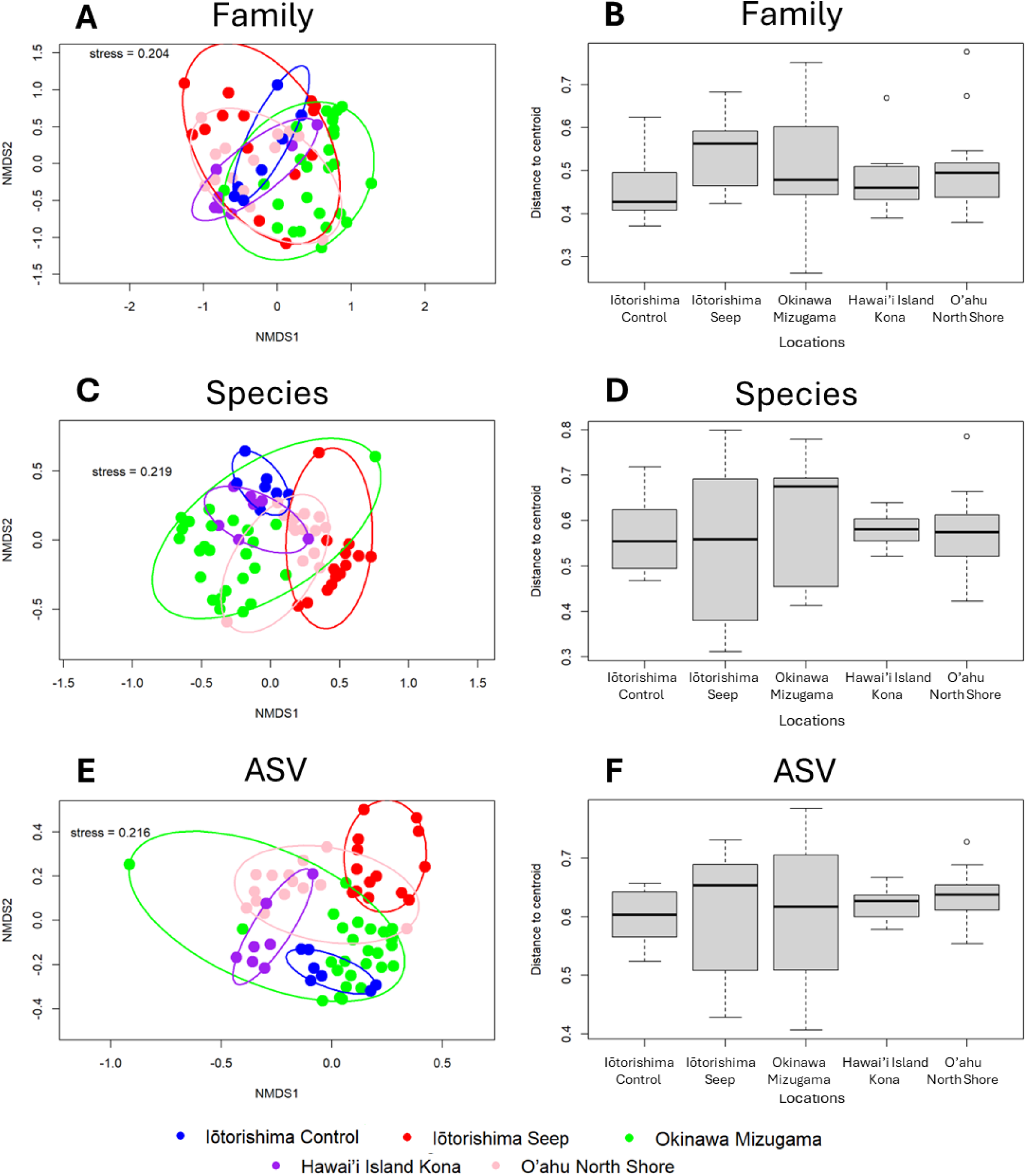
Non-Pac Multidimensional Scaling (A, C, E) and distance to centroid of beta-diversity dispersion (B, D, F) for family-level (A, B), species-level (C, D), and ASV (E, F), hosted by *Palythoa tuberculosa* at different locations. Blue=Iōtorishima Control, Red= Iōtorishima Seep, Green=Okinawa Mizugama, Purple =Hawai’i Island Kona, Pink = O’ahu North Shore.

### Beta diversity - Symbiodiniaceae

There were significant differences among the locations for the Symbiodiniaceae OTU composition (PERMANOVA: R^2^=0.351, F=8.122, p<0.001). Tests of composition dissimilarity between groups showed only Tsukishima Seep *P. tuberculosa* hosted significantly different Symbiodiniaceae compared to all other locations (Table 3, pairwise Adonis: p<0.001). Hawai’i Island Kona specimens had significantly different Symbiodiniaceae composition with the other Pacific locations, except for with O’ahu North Shore (Table 3, pairwise Adonis: R^2^ = 0.152, p = 0.192) Other pairwise comparisons showed no significant differences between locations (Table 3, pairwise Adonis: p>0.192).

**Table 3.** Pairwise PERMANOVA (Pairwise Adonis) conducted on the dissimilarity of Symbiodiniaceae composition for different locations. The lower table represents the adjusted p-values (Holm’s correction) and the upper table represents the R2 values. (statistical significance: ** = 0.01)

|  | Iōtorishima<br>Control | Iōtorishima<br>Seep | Okinawa<br>Mizugama | Hawai'i<br>Island<br>Kona | O'ahu<br>North<br>Shore |
| --- | --- | --- | --- | --- | --- |
| Iōtorishima<br>Control |  | 0.374 | 0.089 | 0.296 | 0.075 |
| Iōtorishima<br>Seep | 0.010** |  | 0.232 | 0.659 | 0.375 |
| Okinawa<br>Mizugama | 0.192 | 0.010** |  | 0.233 | 0.078 |
| Hawai'i Island<br>Kona | 0.010** | 0.010** | 0.010** |  | 0.152 |
| O'ahu<br>North Shore | 0.279 | 0.010** | 0.192 | 0.192 |  |

nMDS (Fig. 4, stress=0.094) showed overlapping Symbiodiniaceae composition between Okinawa Mizugama specimens with other locations. While the Iōtorishima Seep Symbiodiniaceae were completely separated from those of Hawaii main island, there was some similarity with other locations (Supplementary Figure 2). Furthermore, Hawai’i Island Konai specimens had the lowest variation within a site. While agreeing with the beta-dispersal output (Fig. 4), Okinawa Mizugama had the highest median of departure from centroid in diversity with a larger dispersal on nMDS.

**Fig. 4.**
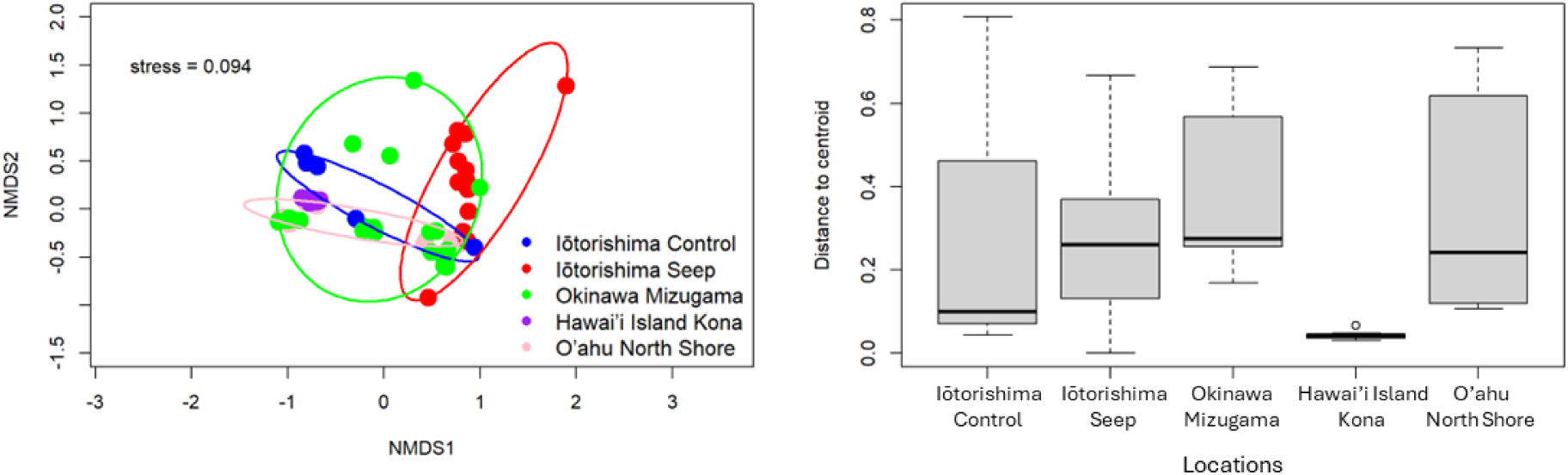
Non-Pac Multidimensional Scaling (left) and distance to centroid of beta-diversity dispersion (right) for Symbiodiniaceae hosted by *Palythoa tuberculosa* at different locations. Purple =Hawai’i Island Kona, Blue=Iōtorishima Control, Red= Iōtorishima Seep, Green=Okinawa Mizugama.

The beta-diversity dispersion of Symbiodiniaceae hosted by *Palythoa tuberculosa* within each location indicated that the median OTU diversity was closely similar in Iōtorishima Seep, Okinawa Mizugama, and Hawai’i Island Kona (Fig. 4). Permutation tests for the size of dispersion relative to the centroid showed significance among the locations (Permutest: Df=4, F=4.6146, p=0.004). Post hoc tests on the diversity dispersion of Symbodiniaceae showed significant differences only for the very low OTU diversity variation of Hawai’i Island Kona compared to other locations (p < 0.05).

SIMPER analyses showed Symbiodiniaceae *Cladocopium* were the main driver of the dissimilarity in Symbiodiniaceae OTU composition, specifically with *Cladocopium* C1, C3, C71 and C1n as the main drivers of differences. Furthermore, C1 and C3 were the dominant drivers of differences between the Iōtorishima Seep and other locations.

### Bacterial community composition

At the phylum level, the bacterial composition was dominated by *Proteobacteria*, irrespective of location (Fig. 5). Unique to the Iōtorishima Seep was the high abundance of *Firmicutes*, the absence of *Cyanobacteria*, and the increased abundance of *Actinobacteria*, comparable to as observed at O’ahu North Shore.

**Fig. 5.**
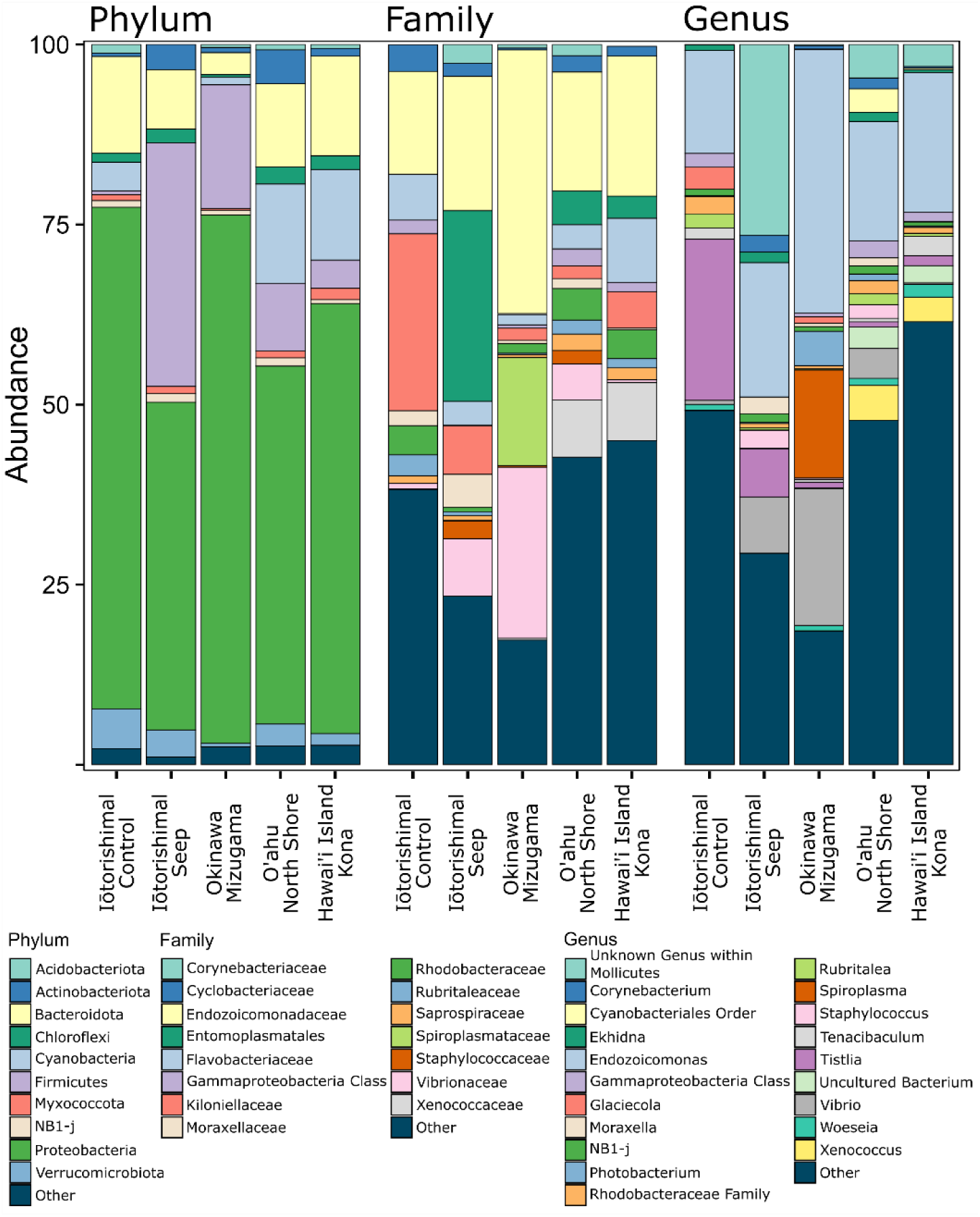
Bar graph showing the bacterial community composition at Phylum, Family, and Genus level for the five investigated sites: Iōtorishima Control, Iōtorishima Seep, Okinawa Mizugama, O’ahu North Shore, and Hawai’i Island Kona.

In terms of family composition, the Iōtorishima Seep location had a high abundance of *Corynebacteriaceae*, *Entomoplasmatales*, *Moraxellaceae*, *Staphylococcaceae* and Vibrionaceae, especially when compared to the Iōtorishima Control site (Fig. 5). On the other hand, the Iōtorishima Control site had a higher relative abundance of *Kiloniellaceae* and *Rhodobacteraceae*.

The bacterial composition at genus level showed a high abundance of *Mollicutes*, *Corynebacterium*, *Moraxella*, and *Vibrio* at the Iōtorishima Seep site (Fig. 5). Interestingly, *Vibrio* was highest at Okinawa Mizugama, which was the most heavily urbanized site, followed by Iōtorishima Seep, and O’ahu North Shore, while being absent in the remaining two sites. Mizugama showed a unique accumulation of *Spiroplasma*. As well, the Iōtorishima Control in particular, but to a lower degree also the Iōtorishima Seep, showed a comparatively high concentration of *Tistlia*, which was uncommon at the other sites.

## Discussion

Our analysis of the Symbiodiniacea and bacterial compositions in the zoantharian *Palythoa tuberculosa* from the high *p*CO_2_ site at Iōtorishima, compared to a control site on the same island, as well as sites from other locations in Okinawa and Hawai’i, revealed both expected and surprising findings. As with any extensive high-throughput sequencing study, some results initially seemed perplexing. However, when interpreted in the context of exisiting knowledge about holobiont Symbiodiniaceae and bacterial compositions, this research provides a strong foundation for future studies and offers important hypotheses on how high *p*CO_2_ levels may impact zooxanthellate anthozoan holobionts.

Firstly, the findings from Symbiodiniaceae analyses in this study warrant discussion. Similar to previous research with *P. tuberculosa* across the Indo-Pacific Ocean (Mizuyama, Masucci, & Reimer, 2018; Reimer, Takishita, & Maruyama, 2006; M. E. Santos et al., 2024; Wee, Kobayashi, & Reimer, 2021), *Palythoa tuberculosa* in our collected specimens from Hawaiʻi was found to be in symbioses with various types of *Cladocopium* (former *Symbiodinium* ‘clade C’; (LaJeunesse et al., 2018). Despite all specimens hosting *Cladocopium*, the variation among *Cladocopium* types was the primary factor driving differences among sites. Specifically, *Cladocopium* C1 and C3-related types were identified to be the main contributors to the distinct Symbiodiniaceae communities at Iōtorishima’s compared to other sites. Additionally, Iōtorishima *P*. *tuberculosa* exhibited lower Symbiodiniaceae diversity than the control sites. These differences in symbiont communities were also clearly depicted in the nMDS analyses, which demonstrated that the high *p*CO2 Symbiodiniaceae communities at Iōtorishima were distinct from those at all other sites, with minimal overlap among ellipses. This is particularly notable considering that the “control” specimens at Iōtorishima were collected less than 200 m away from the high CO_2_ specimens, yet they were more similar to Symbiodiniaceae communities from Oʻahu North Shore and Hawaiʻi Island, over 7,000 km away, as well as to specimens in Okinawa, more than 170 km away. This lower anthozoan community diversity anthozoans thriving in more stressful environments aligns with recent findings showing higher symbiont variability in corals with lower resilience to stressful environments (Howe-Kerr et al., 2020). While it may be premature to conclude that high levels of *p*CO_2_ are the main cause of these differences, it is certainly a strong possibility. Based on our current results and previous findings at the same seep (Wee et al., 2019), we hypothesize that elevated CO_2_ levels may lead to reduced Symbiodiniaceae diversity, at least in some anthozoan hosts like *P*. *tuberculosa*. Similar findings have also been observed in three species of scleractinian corals in Bourake, New Caledonia, which showed more homogenous Symbiodiniaceae communities compared to colonies at control sites (Tanvet et al., 2023). While the mechanism behind this reduction in Symbiodiniaceae under high CO_2_ levels remains to be unclear, is it possible that endosymbiotic Symbiodiniaceae experience higher rates of photosynthesis under lower pH conditions (Tanvet et al., 2023). Our results support the hypothesis that “corals from distinct environments often have unique symbiotic partners that could be crucial to support their survival” (Tanvet et al., 2023). A deeper understanding the environmental preferences and adaptations of various *Cladocopium* types within *P*. *tuberculosa* and other host species could help researchers confirm or refute these hypotheses. Recent research on *P*. *tuberculosa* around Okinawa has confirmed the presence of different lineages with hypothesized preferences for specific environments (Noda, Parkinson, Yang, & Reimer, 2017; Wee et al., 2021), further supporting this idea.

The bacterial findings also reveal interesting patterns. Bacteria, a group that has existed for at least 3.5 billion years, display immense diversity across various phyla, each with its unique biology and a broad range of functions within ecosystems. Although *Palythoa* species have been reported associated with over 30 bacteria families, there was a distinct bacterial composition associated with specimens from the Indo-Pacific or Atlantic oceans (M. E. Santos et al., 2024). This diversity was evident in our results, showing that the *P*. *tuberculosa* holobiont can encompass a highly diverse assemblage of bacteria. For instance, our *P*. *tuberculosa* specimens contained 394 bacterial families in total, with 12 families each contributing ≥ 1% relative abundance. Similarly high levels of bacterial diversity have been observed in many cnidarians, particularly in zooxanthellate anthozoan and scleractinian holobionts (e.g. reviewed in (Pogoreutz et al., 2020). Therefore, the highly diverse bacterial communities from *P*. *tuberculosa* in this study are not unexpected.

However, a closer examination of the bacterial communities reveals several intriguing findings, especially concerning the high *p*CO_2_ site. Notably, the bacterial community richness within *P*. *tuberculosa* from the high *p*CO_2_ site was significantly lower than at any other site. Although family-level analyses did not show many differences across sites, more taxonomically refined analyses at species and ASV levels did reveal distinctions. Additionally, a detailed examination of bacterial community composition, even at the phylum level, indicated potentially important differences (Fig. 5). For instance, at the phylum level, the Iōtorishima high *p*CO_2_ samples had a higher abundance of Firmicutes compared to other sites. At the family level, greater presence of *Entoplasmatales* was observed, and at the genus level there was a higher abundance of an unknown genus within *Mollicutes*. *Mollicutes* genotypes have been widely reported in various marine species (e.g., seastars (Loudon, Park, & Parfrey, 2023) and terrestrial invertebrate (e.g., isopods, (Fraune & Zimmer, 2008). They have also been shown to influence seasonal differences in bacterial communities in rock oysters, and have been implicated in variations in disease resistance (Nguyen et al., 2020).

In anthozoans, the few reports on *Mollicutes* present a different picture compared to our current findings. *Mollicutes* species have been documented in octocorals from the Great Barrier Reef, where their abundances remained stable over time, even before, during, and after a bleaching event (Steinberg, Dafforn, Johnston, & Ainsworth, 2023). Similarly, they were reported as stable in Mediterranean gorgonian octocorals over time (van de Water et al., 2018), leading Steinberg et al. (2023) to hypothesize that *Mollicutes* may play a significant functional role in the microbiome of the host colonies. However, our results contrast with these previous reports. While *Mollicutes* were found in relatively low numbers in both Hawaiʻi and Okinawan *P*. *tuberculosa* specimens, it was almost absent (< 0.01% relative abundance) at the control site at Iōtorishima, but highly abundant (25.6 % relative abundance) at the high *p*CO_2_ site. Although *Mollicutes* may encompass various species or functional groups, and dispite the differences from previous anthozoan studies, our findings support a link between this taxon and pH or CO_2_ levels. It is clear further research is needed into this intriguing matter.

Noteworthy are the changes in the abundance of the family *Vibrionaceae*, particulary the genus *Vibrio*. High abundances of *Vibrio* were observed at Mizugama (15.1% relative abundance), Iōtorishima Seep (6.7% relative abundance), and O’ahu North Shore (6.0% relative abundance), while Iōtorishima control and Hawaiʻi Island exhibited much lower levels (< 1% relative abundance). Some members of *Vibrionaceae* are considered potential opportunistic and pathogenic bacteria, linked to several coral diseases (reviewed in (Mohamed, Ochsenkühn, Kazlak, Moustafa, & Amin, 2023). Their increased presence has been associated with environmental changes (Brumfield et al., 2023), particularly in immune-comprised hosts (e.g., (Rubio-Portillo et al., 2018).

Research on natural analogues has gained momentum in recent years due to growing concerns about the impacts of climate change on marine ecosystems (Leung et al., 2022). On coral reefs, most of the research has focused on scleractinian corals (e.g., (Godefroid, Dupont, Metian, & Hédouin, 2022) and fish (e.g., (Priest et al., 2024). However, an increasingbody of literature suggests that as zooxanthellate scleractinian corals decline, other benthic groups may become dominant on reefs. Reports indicate the potential spread of such “non-coral” reefs due to environmental shifts, involving a wide variety of taxa, including sponges (Bell, Davy, Jones, Taylor, & Webster, 2013), sea anemones (Suggett et al., 2012), corallimorpharians (Tkachenko, Wu, Fang, & Fan, 2007), soft corals (Inoue et al., 2013; Lalas, Jamodiong, & Reimer, 2024), algae (Agostini et al., 2018), and zoantharians (Cruz et al., 2015; Reimer et al., 2023; Reimer, Wee, et al., 2021). Unfortunately, for many of these groups, essential biological data such as growth rates, tolerances, sexual reproductive traits, and even accurate taxonomic identification are often lacking (Otis et al., 2024). This data gap hinders our ability to accurately model or predict which benthic organisms might replace scleractinian corals under various conditions, severely limiting our capacity to forecast the coral reefs. Therefore, further research in this area is urgently needed (Otis et al., 2024).

At the same time, it is important to recognize the significant diversity within each of these understudied groups, and researchers should avoid treating them as a single entity (Cannon et al., 2023; Otis et al., 2024). In this study, we focused on *P. tuberculosa*, a common and widely distributed Indo-Pacific species (Hibino, Todd, Ashworth, Obuchi, & Reimer, 2013). Recent research suggests that this species is flexible in terms of its heterotrophic performance (M. E. A. Santos, Baker, Conti-Jerpe, & Reimer, 2021) and symbiont composition, capable of hosting different *Cladocopium* lineages depending on local conditions, which allows it to thrive in a wide range of habitats (Noda et al., 2017; Wee et al., 2021). However, it would be unwise to assume that all zoantharians, including those common on coral reefs (e.g., genus *Zoanthus*), will behave similarly, even under low pH conditions. There is also the possibility that *P. tuberculosa* represents multiple closely related lineages (Dudoit, Santos, Reimer, & Toonen, 2022).

Finally, there is an even greater scarcity of data on the holobionts of these groups. In fact, there are very few published holobiont datasets for both Symbiodiniaceae and bacteria in *P. tuberculosa* (with the exception of (Paulino, Broetto, Pylro, & Landell, 2017) and none for natural analogues. Given the abundance of unique “understudied” taxa at natural analogues, it is crucial to urgently aquire data on these holobionts, along with the basic biological information mentioned eariler. Simultaneously, efforts should be made to gather more comprehensive holobiont datasets at higher resolutions than are currently available, covering a broad geographic range.

In conclusion, marginal environments such as the high *p*CO_2_ reef at Iōtoroshima have a distinct diversity and composition of Symbiodiniaceae and bacteria associated with *P. tuberculosa*. Our findings indicate that different components of the holobiont are affected in distinct ways and detecting these variations often requires powerful, high-resolution methods and comparisons across both narrow and wide scales. To fully understand and better predict the impacts of climate change on coral reef organisms, it is essential to investigate the various partners involved.

## Acknowledgements

We thank the captain and crew of the *Yosemiya III*, and cruise members Y. Ide (Oceanic Planning Corp.), H. Takamiyagi (OIST), and H. Kayanne (U. Tokyo) for their support and advice at Iwotorishima. We also thank Akito Shima for making the mycoplasma trees

## Funding

This project contributes towards the International CO_2_ Natural Analogues (ICONA) Network. JDR and TR were partially funded by the Japan Society for the Promotion of Science (JSPS) Core-to-Core Program (Grant Number: JPJSCCA20210006). Work at Iōtorishima was supported by an OIST KICKS grant entitled “Japanese volcanic CO2 vents—natural laboratories to study the behavior and adaptation of marine organisms to acidifying oceans” to TR, HK, and JDR. We thank the captain and crew of the *Yosemiya III*, and cruise member Y. Ide (Oceanic Planning Corp.) for their support and advice at Iōtorishima. H. Takamiyagi (OIST) provided support at Iōtorishima. JDR was also partially supported by a JSPS Grant-in-Aid for Transformative Research Areas entitled Environmental, ecological, and genetic observations of coral reef Symbiodiniaceae-host holobiont symbioses (23H03821). HK, MM, GYS and AI were supported by Environmentally-conscious Developments and Technologies (E-code) research laboratory at the National Institute of Advanced Industrial Science and Technology (AIST). HK was also supported in part by JSPS KAKENHI grant number 23KJ2206 from the Japan Society for the Promotion of Science.

## Supplementary Material

**See attached PDF file for supplemental Figure 1.**

**Supplemental Figure 1** The tree was generated using *Mollicutes* ASV (identified as Candidatus *Hepatoplasma* by DADA2) and published data (Vohsen et al., 2024). The tree was constructed using MAFFT (v7.525) with the following command: “mafft -- adjustdirection --globalpair --maxiterate 1000” and IQ-TREE2 (v2.2.2.5) with the GTR+F+R3 model and 1,000 bootstrap iterations.

**Supplemental Figure 2.**
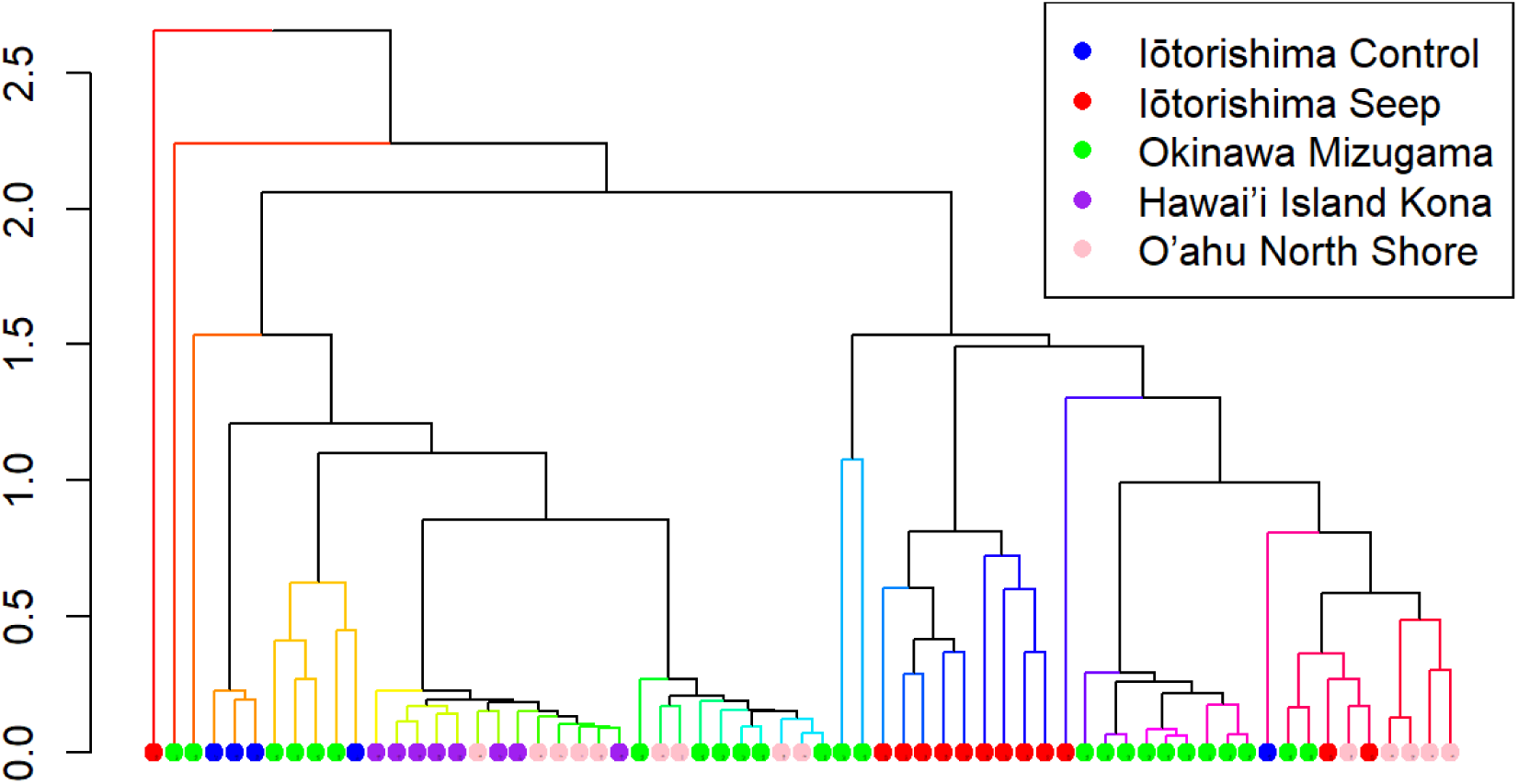
Similarity Profiling (SIMPROF) dendrogram representing the composition of Symbiodiniaceae by *Palythoa tuberculosa* at different Locations (Iōtorishima Control=Blue, Iōtorishima Seep=Red, Okinawa Mizugama =Green, Hawai’i Island Kona= Purple, O’ahu North Shore = Pink).

## Notes

### Competing Interest Statement

The authors have declared no competing interest.

