## Supplementary figures and images for "Shifts in Coral Reef Holobiont Communities in the High-CO_2_ Marine Environment of Iōtorishima Island"

### Supplemental Figure 1

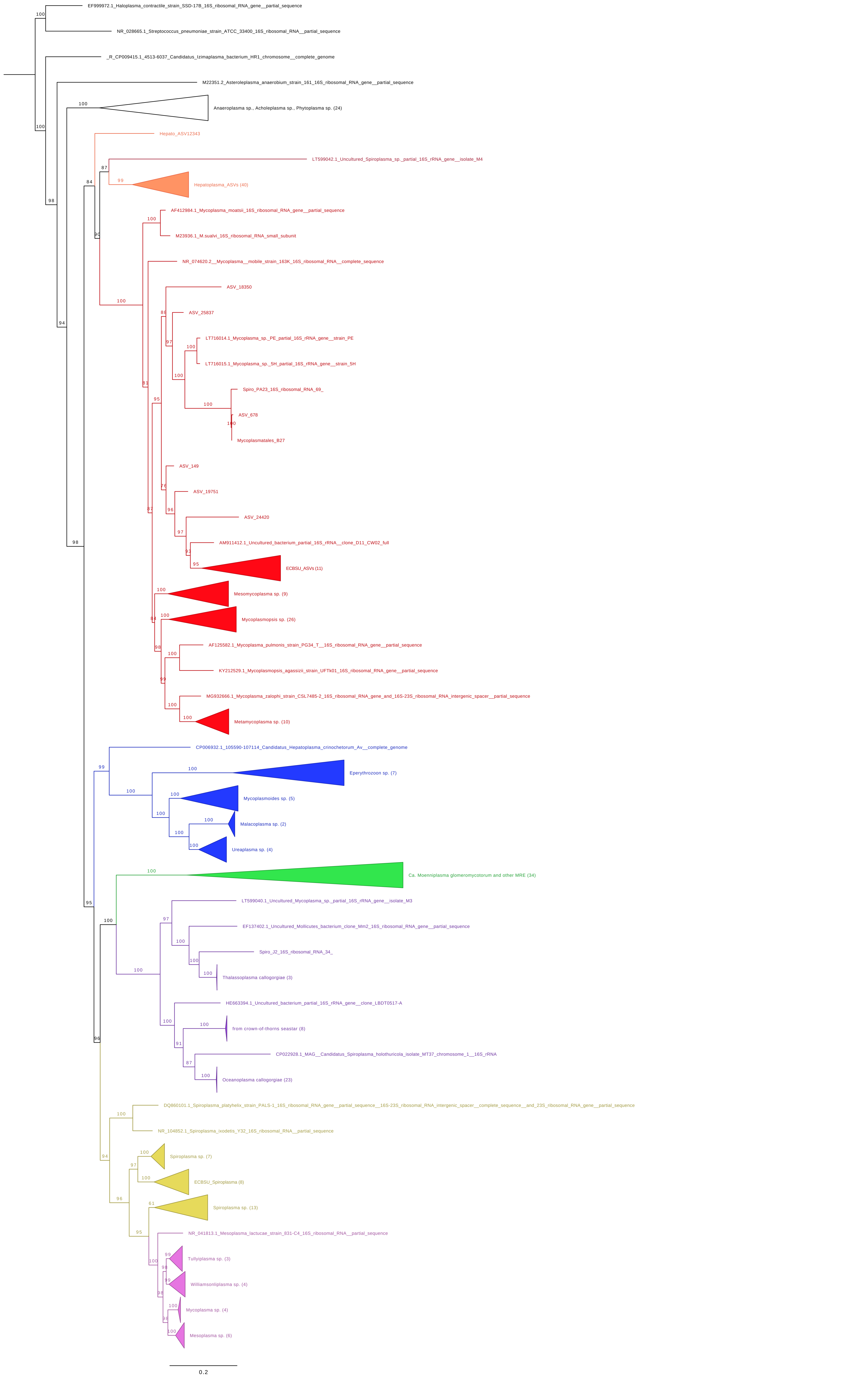
